# A novel Vector-Symbolic Architecture for graph encoding and its application to viral pangenome-based species classification

**DOI:** 10.1101/2025.09.08.674958

**Authors:** Fabio Cumbo, Kabir Dhillon, Jayadev Joshi, Davide Chicco, Sercan Aygun, Daniel Blankenberg

## Abstract

Viral species classification is crucial for understanding viral evolution, epidemiology, and developing effective diagnostics and treatments. Traditional methods often rely on sequence similarity, which can be challenging for rapidly evolving viruses. Pangenomes, offering a comprehensive representation of species’ genomic diversity, provide a richer perspective, but their analysis often requires advanced computational methods. We investigate the use of Hyperdimensional Computing (HDC), also known as Vector-Symbolic Architecture (VSA), an emerging computing paradigm that relies on vectors in high-dimensional spaces to encode a multi-species viral pangenome.

We develop a new method for encoding graph-structured viral pangenomes using high-dimensional vectors. Pangenomes are represented as weighted *de Bruijn* graphs constructed using sequences of consecutive k-mers from the genomes, while information about the genome species (their class) is encoded as specific weights on the edges of the graph. The weighted *de Bruijn* graph representation is encoded into a single high-dimensional vector. We tested three classification strategies: a flat model at the species level, a flat model at the genus level, and a two-step hierarchical model.

We applied our method to a pangenome comprising 542 viral species from NCBI GenBank. Our results reveal a complex relationship between model architecture and classification accuracy. The flat species-level model achieved the highest accuracy, correctly classifying 87.08% of test genomes. Counter-intuitively, simplifying the problem to the genus level or using a hierarchical approach degraded performance, with accuracies dropping to 60.51% and 33.57% respectively. These outcomes highlight critical challenges in alignment-free classification, such as signal dilution in overly broad taxonomic groups and error propagation in multi-step models. The model’s reconstruction rate proved to be a reliable measure of confidence, rather than a direct predictor of correctness.

This novel approach offers a promising new direction for viral classification, not only for its predictive power but its ability to reveal underlying challenges in genomic taxonomy.

## 1. INTRODUCTION

Viral species classification is a cornerstone of modern virology, essential for understanding viral evolution, tracking epidemiology, and developing diagnostics and therapeutics [1]. However, classifying viruses presents profound challenges rooted in their biology. Their rapid mutation rates, high frequency of genetic recombination, and vast diversity mean that traditional classification methods, which often depend on sequence similarity of conserved marker genes, are frequently inadequate [2]. Unlike cellular life, viruses lack a universally conserved gene analogous to the 16S rRNA gene, making the construction of a comprehensive viral tree of life a complex, ongoing endeavor [3–6]. This biological reality necessitates the development of advanced, scalable, and accurate classification methods that can handle the ever-expanding volume of viral genomic data.

To better capture the full spectrum of viral genetic diversity, the field has increasingly adopted a pangenomic perspective [7–10]. A pangenome represents the entire gene repertoire of a species, encompassing the core genes (gene shared by all strains) and the accessory genes (genes present in only a subset of strains) [11,12]. This approach provides a more holistic view of a species’ adaptive potential, revealing insights into functional diversity, mechanisms of virulence, and evolutionary trajectories. Pangenomes are often visualized as complex graph structures, where nodes represent genetic sequences and edges denote their adjacency. These pangenome graphs are powerful representations that can capture large-scale structural variations, such as insertions, deletion, and rearrangements, which are often missed by linear sequence alignment [13].

While pangenome graphs offer a rich representation, their analysis introduces significant computational challenges. Constructing pangenomes with established gene-based methods often requires computationally expensive algorithms. For instance, creating a core-gene phylogeny, a key output, depends on multiple sequence alignment, which can become a major bottleneck when dealing with thousands of genomes [14]. As sequencing projects continue to deposit massive datasets into public repositories, the need for more efficient and scalable methods for pangenome analysis has become critical. The challenge lies in developing a framework that can both manage this scale and accurately model the complex relationships inherent in pangenomic data.

In this context, Hyperdimensional Computing (HDC, also known as Vector-Symbolic Architectures (VSA), emerges as a promising computational paradigm inspired by principles of neural computation [15–17]. HDC operates on high-dimensional vectors (often referred to as hypervectors), using a defined set of mathematical operations to represent and manipulate complex data structures and relationships between information. This framework offers several key advantages for bioinformatics applications: its distributed nature makes it inherently robust to noise and errors (such as sequencing errors or minor genomic variations), its reliance on simple, parallelizable operations addresses computational bottlenecks, and its capacity for one-shot learning is well-suited for biological classification tasks [18,19].

The unique properties of HDC have led to its successful application across a range of bioinformatics problems. In genomics, HDC has been used to develop fast, alignment-free methods for sequence comparison and analysis [20–24]. It has also proven effective in classifying complex biological data, from DNA methylation patterns in cancer [25] to microbial abundance profiles in metagenomics [26], and has been applied to challenges in cheminformatics, such as drug discovery and classification [27–29] among other applications to complex problems in the life sciences [18,19]. The ability of HDC to create compact yet descriptive representations of complex data makes it an ideal candidate for tackling the challenge of pangenome analysis, an area where its potential has not been explored so far.

This work introduces a novel VSA-based framework for viral species classification that directly leverages the power of pangenome graphs. We model the viral pangenome as a weighted *de Bruijn* graph [30] where nodes represent k-mers and edges are colored with hypervectors corresponding to the species in which they appear. This complex graph is then encoded into a single, holistic hypervector. This approach allows us to efficiently query the pangenome to determine the species of a new viral genome.

Our main contributions are:

1. a novel method for encoding large, weighted/colored *de Bruijn* graphs into a single, high-dimensional vector representation;
2. the application and validation of this method for the complex task of viral species classification;
3. a demonstration of a scalable and hierarchical framework that improves classification accuracy by creating specialized, genus-specific models.
4. experimental proof of linear relation between graph reconstruction and the corresponding accuracy.

The method leverages the right information contained within pangenomes and the computational efficiency of VSAs to enable accurate viral species classification at scale, offering a powerful new tool for computational virology.

## 2. MATERIALS AND METHODS

Here we present the data we used for our analysis in addition to an introduction to the theoretical foundations of the Hyperdimensional Computing paradigm together with our graph-based encoding technique.

### 2.1. Data acquisition and preprocessing

We retrieved all the viral reference genomes from the NCBI GenBank database [31] and randomly selected two genomes for each of the 542 species, for a total of 1,084 viral genomes belonging to 218 genera, 61 families, 37 orders, 24 classes and 14 phyla. Our definition of reference genomes was expanded to include any GenBank assembly not excluded from NCBI RefSeq for the following reasons, which we group into two main categories:

- <u>Issues with source material or provenance</u>: (i) derived from single cell – the assembly was generated from single-cell amplified material, which can impact sequence accuracy; (ii) non-standard type material – the assembly was constructed from sequences designed as various type materials;
- <u>Issues with metadata or project context</u>: (i) incomplete taxonomic information – the assembly lacked a specified strain identifier or had an undefined genus in its lineage; (ii) part of a large-scale project – the assembly was flagged as belonging to a large project with over 100 isolates of the same species, which often have different data-release and annotation standards.

Accession numbers of the reference genomes considered in our analysis are provided in Supplementary Table S1 alongside with their taxonomic label as provided by NCBI GenBank.

It is worth noting that we focused on viral species with at least 2 reference genomes in NCBI GenBank, also performing a quality control with CheckV [32] by filtering out low- and medium-quality genomes with a completeness <90% and a contamination >5%. Then, we randomly selected 2 genomes per species among those that survived the quality filtering.

Since our method is k-mer-based, we first fragmented all the selected genome sequences into overlapping k-mers of length 9 over a sliding window with a single nucleotide step, by also maintaining their order of occurrence in the original sequence. We determined k-mer length empirically by computing the Cumulative Relative Entropy, the Average Number of Common Features, and the Observed Common Features for each *k* in the closed interval [6 − 20] with *Kitsune* [33], a Python package for the estimation of the optimal k-mer size for a specific set of genomes.

### 2.2. Hyperdimensional Computing and the MAP model

HDC is a promising computing paradigm inspired by the way the human brain works in encoding information [15,16,19]. It uses random binary or bipolar high-dimensional vectors, often called hypervectors, as atomic computing data. In particular, to represent and manipulate symbolic information, HDC excels by leveraging orthogonal representations with vector sizes typically ranging from *D* = 100 to *D* = 1000 bits, with *D* always much greater than the number of atomic information *N* to be represented in the high-dimensional space (a *D*-dimensional space guarantees the existence of up to *D* orthogonal vectors in the same space; *D*≫*N* to increase the chance of orthogonality in case of randomly generated vectors). Unlike conventional machine learning and neural networks, HDC targets single-pass, error- and backpropagation-free learning methods, building on bio-inspired concepts such as associative and item memory, naturally offering a series of advantages like:

- <u>Robustness to noise and error</u>: HDC uses high-dimensional vectors, distributing information across all dimensions. Scalars or symbols are represented using long-dimensional vectors, where the cumulative counts of binary values and their ratios encode the information. However, this encoding is fundamentally different from traditional binary radix-2 representation, as it does not assign significance to any specific bit. This inherent redundancy makes HDC highly tolerant to noise and errors, unlike conventional computing, where a single bit flip (e.g., the most significant or sign bit) can drastically alter the outcome. This robustness is crucial for real-world applications where data can be noisy or incomplete [34,35];
- <u>Computational efficiency</u>: HDC operations are simple and highly parallelizable, leading to fast computation, especially for tasks like similarity search in high-dimensional spaces. Algebraic operations (e.g., multiplication and addition) can be implemented with simple logic during encoding, which reduces computational complexity, particularly in application-specific hardware designs [36,37];
- <u>One-shot learning</u>: HDC can often learn from single examples, unlike deep learning, which requires extensive training, large datasets, batching, and randomization. In contrast, HDC depends only loosely on repeated data input and requires little to no heavy feature engineering. This ability to generalize from limited data makes it suitable for situations where training data is scarce or expensive [38–40];
- <u>Power and energy efficiency</u>: due to its simple and parallelizable operations, HDC can be implemented in power- and energy-efficient hardware. Recent advances in vector design techniques further improve this efficiency: shorter vectors enable lightweight architectures with lower latency, making HDC particularly advantageous in energy-constrained scenarios [41,42];
- <u>Interpretability and transparency</u>: the symbolic nature of HDC can make its operations more interpretable than those of deep learning models, which are often considered “black boxes” [19,43];
- <u>Scalability</u>: following the previous point about the computational efficiency, HDC can perfectly scale to large datasets. Through encoding, repeated or complex data can be compressed into one-dimensional vectors, which enables efficient representation and supports scalability across diverse data sizes [44,45];
- <u>Flexibility and generality</u>: HDC can represent and manipulate a wide range of data types and structures [46], making it adaptable to diverse domains and applications [47]. This flexibility is evident in its use for various bioinformatics tasks [18,19] as well as other applications like natural language processing [48], artificial intelligence [49,50], robotics [51], among other scientific domains.

The reason hypervectors are typically large in size is because of their random nature. The higher the dimensionality *D*, the higher the chances that two randomly generated vectors are orthogonal or quasi-orthogonal in the same space (with a cosine-similarity close to 0).

All these characteristics make HDC well-suited for handling large and complex datasets, particularly in bioinformatics and other data-intensive fields.

HDC operates on hypervectors using a small but efficient set of arithmetic operations, i.e., *multiplication, addition*, and *permutation*, that together compose the MAP (Multiply-Add-Permute) model:

- <u>Multiplication</u> (a.k.a. binding): this operation binds two hypervectors together to create a new hypervector that represents the association or relationship between the original two. The resulting hypervector is dissimilar to both input vectors. This operation is crucial for representing complex structures and relationships. The specific implementation of multiplication can vary depending on the chosen vector space, i.e., it could be implemented as the component-wise XOR in case of random binary vectors (with 0s and 1s), or component-wise multiplication in case of random bipolar vectors (with −1s and +1s). Using binary vectors and XOR in hardware simplifies the algebraic operation in a cost-effective manner;
- <u>Addition</u> (a.k.a. bundling): this operation bundles multiple hypervectors together to create a new hypervector that represents the set or combination of the input vectors. The resulting hypervector is similar to all input vectors. It is implemented as the component-wise majority rule in case of binary vectors, or component-wise addition in case of bipolar vectors, which is typically followed by a thresholding step (e.g., majority voting) to keep the result within the same vector space. This operation is essential for representing sets, sequences, and other composite structures.
- <u>Permutation</u>: it shifts the elements of a hypervector by a certain number. This helps keep positional or temporal features based on their order. It can represent sequences and temporal relationships between concepts (e.g., a permuted hypervector might represent the next item in a sequence, such as consecutive letters in a language processing application).

The combination of these three operations allows for complex computations and manipulations of high-dimensional representations, enabling HDC systems to perform various tasks, including classification, analogy reasoning, and language processing.

### 2.3. Graph encoding

Here, we present a HDC approach to encoding directed, weighted graphs, which we adopt and adapt from a previous study by Prathyush Poduval *et al*. [52]. To encode a directed weighted graph *G* = (*V, E, W*) with *V* the set of nodes, *E* the set of edges, and *W* the set of weights, we assign a random bipolar hypervector *H* ∈ {− 1, + 1}^*D*^ to each node *ν*_*i*_ ∈ *V*. These vectors are quasi-orthogonal due to their random nature as we previously explained.

#### 2.3.1. Encoding directed connections

For each node *ν* _*i*_, we define a node memory vector *M*_*i*_ to capture all the directed outgoing connections from *ν*_*i*_ to its neighboring nodes. In directed graphs, edge directionality must be preserved. In order to do so, we introduce a permutation operator ρ, which rotates a hypervector by a fixed number of positions. This is applied to the memory component to break the symmetry in the binding operation. Specifically, an edge from node *i* to node *j* is encoded as *H*_*i*_ ⊗ ρ(*M*_*i*_), where ⊗ denotes binding, and ρ applies a permutation that ensures *H*_*i*_ ⊗ ρ(*M*_*i*_) ≠ *H*_*j*_ ⊗ ρ(*M*_*j*_), even if *i* = *j* and *M*_*i*_ = *M*_*j*_, thus capturing the directionality.

The global graph memory *G* is obtained by summing the bound representations of each node and its directed memory: 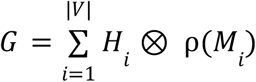. This bundled hypervector encodes the entire directed graph in a single, holistic representation.

#### 2.3.2. Encoding edge weights

In order to encode weights, we employ a deterministic transformation to map continuous weights *w*_*ij*_ ∈ [0, 1) into hypervectors 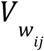. This transformation flips a proportion of components in a base hypervector *V* to create a weighted vector such that 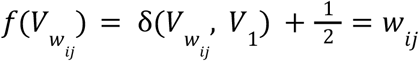. This encoding is deterministic and positionally consistent: the components from index [*w*_*ij*_ · *D*] to *D* in *V*_1_ are flipped. This results in hypervectors for similar weights being more correlated to each other than to those representing distant weights. For a node *i*, its outgoing weighted connections are thus encoded into its memory as 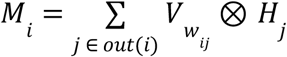, where *out*(*i*) denotes the set of all nodes reachable from *i* via a directed edge. This ensures that not only the connectivity but also the weight of each edge is holographically embedded. The overall encoding procedure for building the graph model is outlined in Algorithm 1.

##### Algorithm 1

VSA-based encoding of a weighted, directed graph. The algorithm outlines the procedure for converting a graph into a single high-dimensional vector *G*. First, a unique, random hypervector *H*_*i*_ is assigned to each node *ν* _*i*_ (lines 2–5). The core of the procedure iterates through each node to compute its local memory. For each node *ν* _*i*_, its memory *M*_*i*_ is created by bundling the representations of all its outgoing connections (lines 11–16). Each connection is encoded by binding the hypervector of the target node *ν* _*j*_ with a hypervector 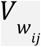 representing the edge’s weight. To preserve directionality, this memory *M*_*i*_ is permuted using a rotation operation ρ (line 18). The node’s own hypervector *H*_*i*_ is then bound to its permuted memory 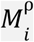 (lines 20–21). Finally, this local representation is bundled into the global graph hypervector *G* (line 23). The final vector holographically represents the entire graph’s topology and weights.

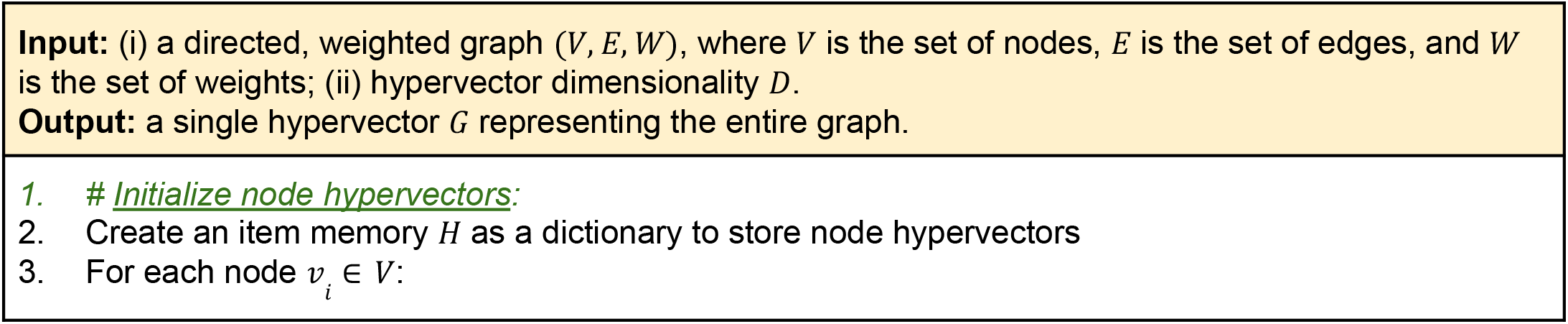

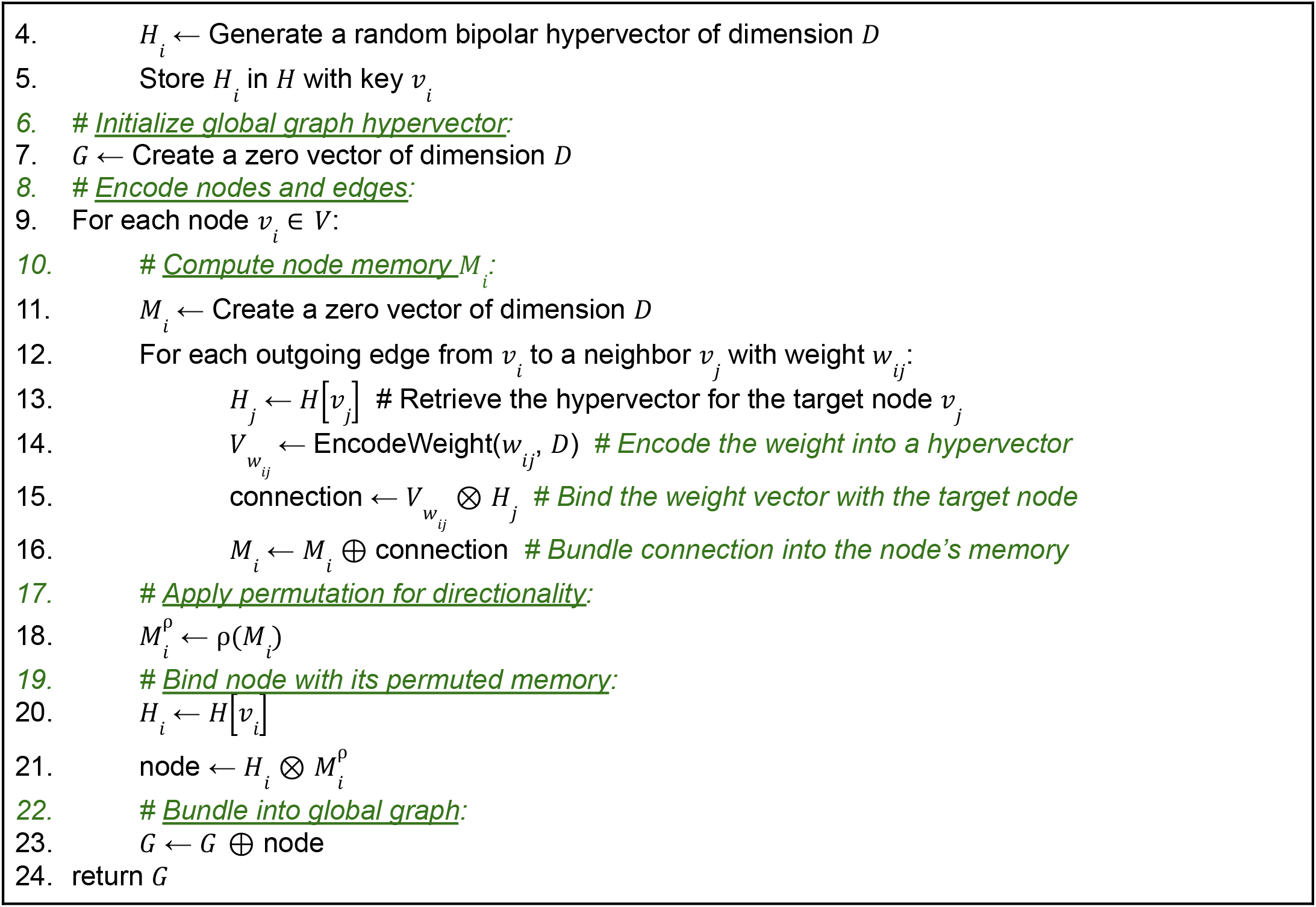

#### 2.3.3. Reconstruction

Given the global memory *G*, local structures can be approximately reconstructed by unbinding and reversing the permutation: 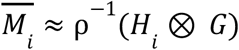. This estimated 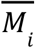 may contain noise due to imperfect orthogonality and vector collisions. To improve recovery, we employ an iterative error mitigation technique called retraining that recomputes each node’s memory by implementing a supervised learning rule.

### 2.4. Retraining and inference

The *retraining* process addresses the specific challenge of a multi-label graph where a single edge can be associated with multiple valid weights (i.e., species labels). A naive correction that subtracts a mispredicted weight is problematic, as that weight might be a legitimate label for that edge in another context. To avoid corrupting valid data, we implement a *one-side retraining strategy* that only corrects for false negatives.

This process is as follows: First, we compute the error rate for the graph model, defined as the number of misclassified edges over the total number of training edges. If the error rate is greater than zero, we begin an iterative correction loop. For each misclassified edge, the model has failed to identify the true weight, indicating its signal is too weak. We therefore reinforce this correct signal by additively bundling its vector representation (the target node bound with the true weight) into the source node’s memory. This procedure only ever adds correct information, ensuring that no valid signals are ever removed or diminished.

This additive update is performed for all misclassified edges in the training set. Afterward, the error rate is recomputed. The entire process is repeated iteratively until the error rate converges to zero, a maximum number of retraining iterations is reached, or the process stops early if the error rate at the current iteration is greater than the rate at the previous one. This leads to a progressive refinement of the model’s memory by strengthening the correct signals until they are reliably detectable.

The process of retraining is summarized in Algorithm 2.

#### Algorithm 2

Iterative retraining of the VSA graph model. This algorithm describes the supervised, one-sided learning process used to refine the graph hypervector *G*. The goal is to minimize classification errors on the training set by selectively reinforcing correct signals without corrupting existing valid information, a crucial step for multi-label graphs. The process begins by calculating the initial error rate, defined as the proportion of misclassified edges (Lines 8–14). An edge (*ν*_*i*_, *ν*_*j*_) is considered misclassified if the model’s prediction for its weight *w*_*pred*_ does not match the true weight *w*_*true*_. The algorithm then enters an iterative loop that continues until the error rate is zero, stops improving, or a maximum number of iterations is reached. The core of the algorithm is the correction step (Lines 20–27). For every edge identified as a false negative, the algorithm computes a reinforced signal. This signal is the vector representation of the correct connection that was too weak to be detected. This corrective signal is then additively applied to the global graph model. This one-sided approach ensures that only correct information is strengthened, preventing the accidental removal of other valid weights that may exist on the same edge. The process repeats, progressively refining the model’s accuracy by amplifying the correct signals until they are reliably detectable.

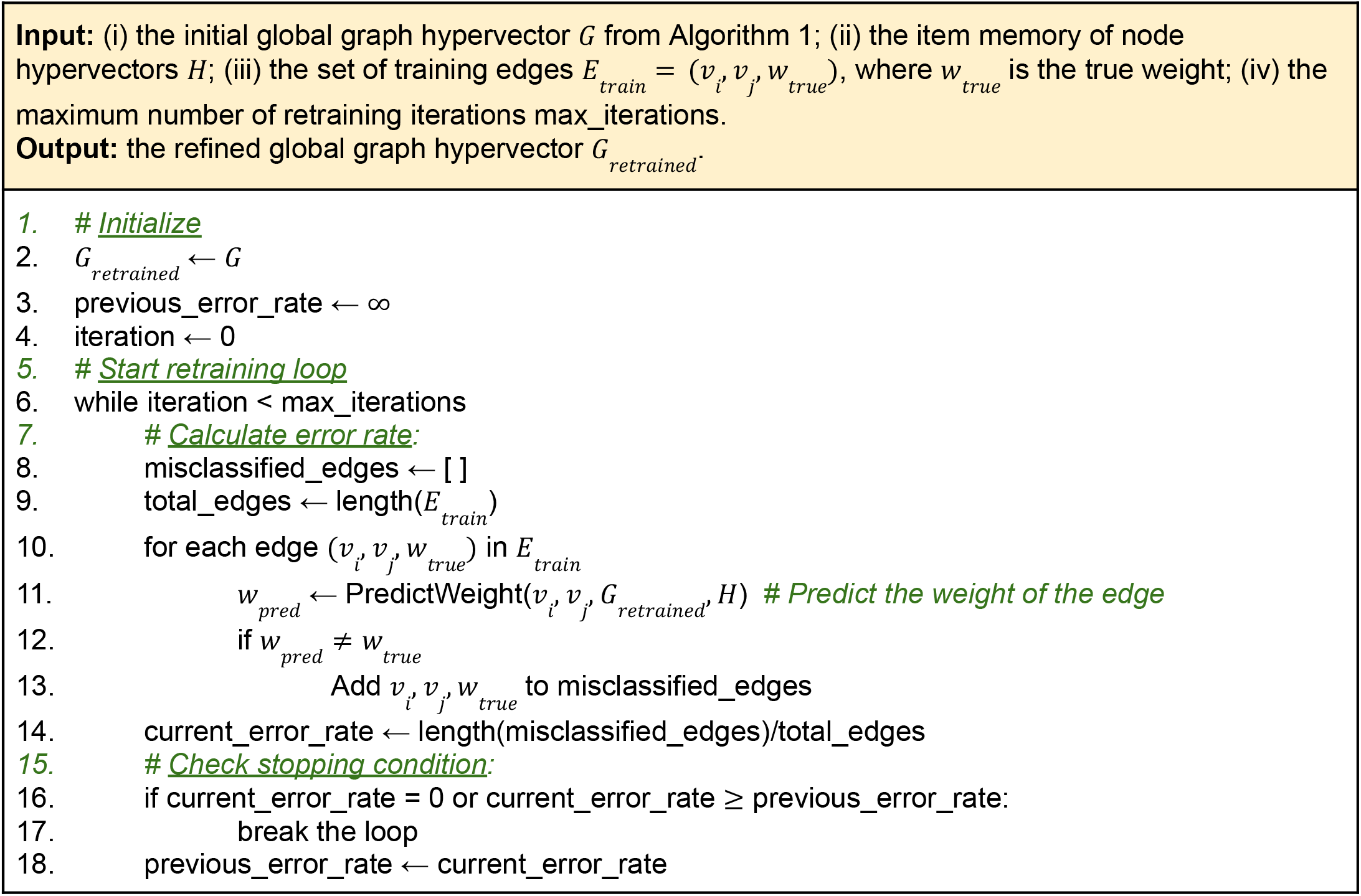

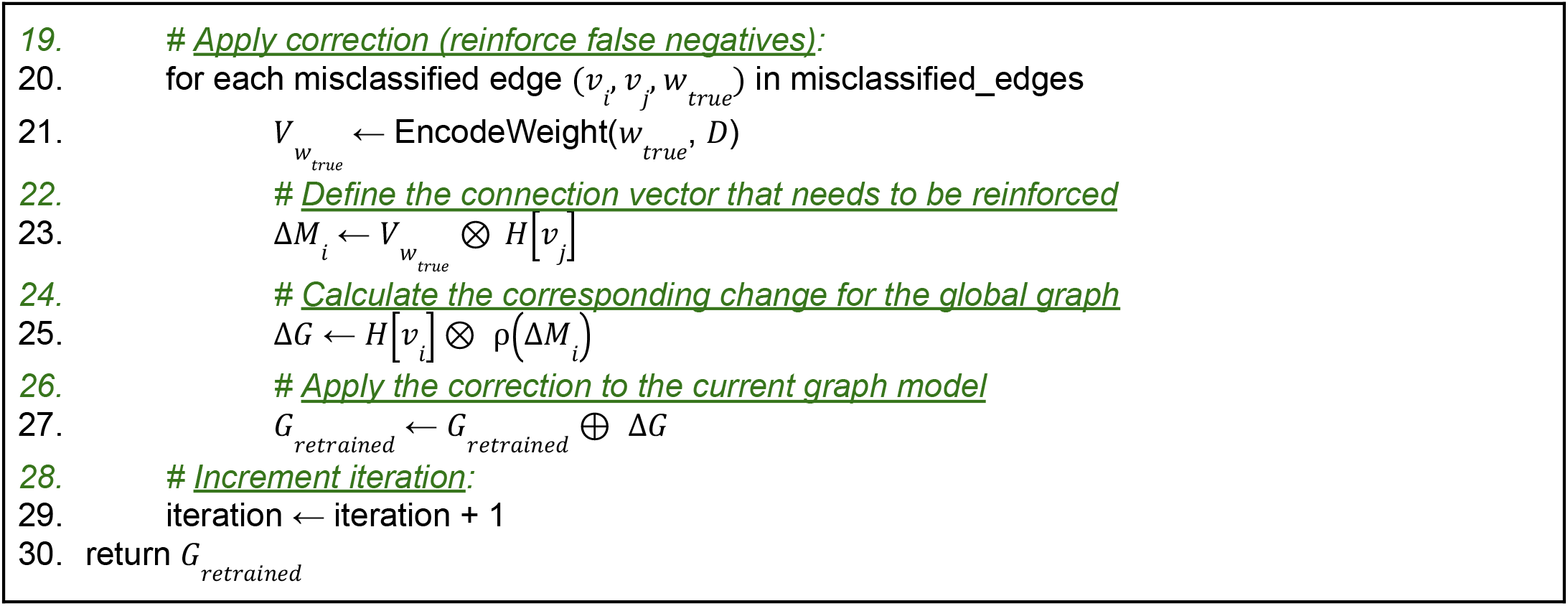

The complete methodology, from the initial graph construction using Algorithm 1, through model refinement with Algorithm 2, to the final prediction on new data, is illustrated as a comprehensive workflow in Figure 1.

**Figure 1.**
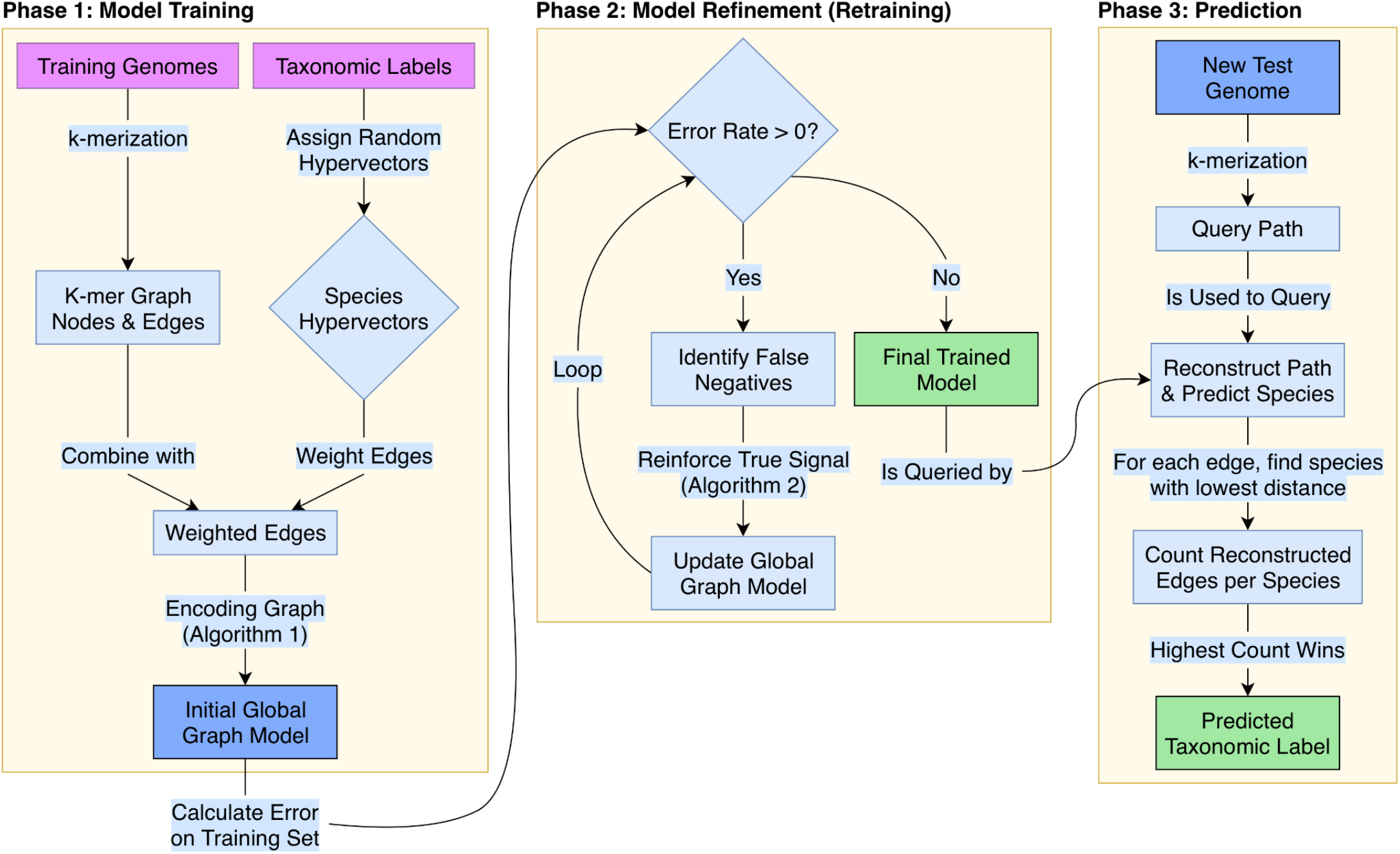
Overview of the VSA-based pangenome analysis workflow. The flowchart illustrates the three main phases of the methodology. <u>Phase 1: Model Training</u> – training genomes and their corresponding taxonomic labels are used to construct an initial weighted *de Bruijn* graph. The labels are converted into unique species hypervectors which color the graph’s edges. This entire structure is then encoded into a single global hypervector following the procedure in Algorithm 1; <u>Phase 2: Model Refinement (Retraining)</u>– the initial model undergoes an iterative retraining loop where false negatives from the training set are identified. The signals for these correct but missed connections are then reinforced in the model as described in Algorithm 2, producing a final, optimized model; <u>Phase 3: Prediction</u> – a new, unseen test genome is decomposed into a query path of k-mers. This path is used to probe the final model, and the system predicts the species label that allows for the highest number of edges to be successfully reconstructed from the graph.

## 3. RESULTS

This section presents the findings of our study on using the HDC-based graph encoding for representing viral pangenomes with the aim to assess the efficacy and advantages of this novel approach in capturing the complex relationships and genomic diversity of all the viral species known so far.

### 3.1. Viral pangenome as weighted *de Bruijn* graph

Pangenomes are typically represented as graphs where nodes correspond to genomic elements and edges represent their adjacency or shared sequences [30]. Here, we constructed a *de Bruijn* graph to represent the collective genetic information of the set of 1,084 viral genomes. A *de Bruijn* graph is a directed graph where nodes represent sequences of a fixed length, called k-mers, and directed edges connect two k-mers if the suffix of the first k-mer matches the prefix of the second k-mer, indicating an overlap of length *k* − 1. For our purposes, k-mers were extracted from the complete genome sequences of the viral species in our dataset. Specifically, for each viral genome, we slid a window of size *k* along its entire length, extracting every possible subsequence of length *k*. This process generates a comprehensive set of k-mers that collectively capture the genomic content and its variations. The choice of *k* is crucial, as a smaller *k* leads to more connections and potentially a more fragmented graph, while a larger *k* can resolve more complex regions but may lead to a sparser graph. Here, we chose *k* = 9 as previously described, and k-mers are extracted sequentially and kept in the order of their appearance in the original sequence.

The graph structure is defined by k-mer adjacency. A directed edge connects two k-mer nodes if they appear consecutively in a viral genome. Additionally, in order to encode the origin of each connection, we assign a unique weight to each of the 542 viral species. Thus, edges are labeled with the specific weight representing different taxonomic labels.

### 3.2. K-mers space reduction with minimizers

To manage the computational complexity associated with handling all possible k-mers, especially in large pangenomes, we employed the concept of minimizers [53]. A minimizer is a subsequence of length *k* that is chosen as a representative of a set of k-mers within a specific window of length *w*, with *w* > *k*. The representative k-mer is established by selecting the first k-mer in the set of alphabetically ordered k-mers of length *k* within the window of length *w*. The process is iteratively repeated by scrolling a window of the same length *w* nucleotide-by-nucleotide over the whole genome. This approach significantly reduces the number of k-mers that need to be processed and stored while still retaining sufficient genomic information to accurately represent the input sequences. Here, we extracted the list of minimizers in the order they appear in the original sequences by choosing a window of length *w* = 50, while maintaining a k-mer size *k* = 9.

### 3.3. Problem formulation and classification results

We designed a VSA to encode viral genetic information into a graph data structure with the aim of assigning species labels to given viral genome sequences. Formally, given a test genome *g*_*test*_ and a set of *C* = 542 possible species classes, the goal is to predict the correct label *c*_*pred*_ ∈ {*S*_1_, *S*_2_,…, *S*_*C*_}. Our approach frames this as a query operation against a holistic, high-dimensional knowledge base. The pangenome, constructed from a training set of genomes, is encoded into a single hypervector *G* that acts as an associative memory. The classification of a new genome is then treated as a process of querying this memory to retrieve the most closely associated species label. The query itself is a hypervector (as in item memory) derived from the test genome’s sequence of minimizers. The success of this approach mostly relies on the ability of the VSA model to (i) encode the complex relational structure of the multi-species pangenome graph into a single vector *G*, and (ii) faithfully reconstruct the species information (the edge weights) when presented with a query path corresponding to a test genome.

In order to test our model’s classification power, we initially partitioned our set of viral genomes into a training and test set with 1 genome per species (i.e., 542 genomes) in each partition. The set of consecutive minimizers in the training set compose the edges that we encoded and collapsed into a single vector representation of the pangenome graph as previously explained. On the other hand, the classification of a test genome proceeds through the following steps:

1. <u>Query generation</u>: the test genome is first decomposed into its ordered sequence of minimizers (using the same *k* = 9 and *w* = 50 parameters) to form a graph representation of the test genome;
2. <u>Memory probing</u>: each of the edges that form the graph representation of the test genome is used to probe the main pangenome hypervector *G*. Given an edge < *A, B, Sp* > where *A* is the source node, *B* is the target node, and *Sp* is a viral species, this is achieved by unbinding the vector representation of the source node *A* from the hypervector *G* to retrieve a new vector (the node memory) that contains information about its neighbors in the pangenome graph;
3. <u>Label prediction</u>: to predict the species of a test genome *g*, we determine which species’ weight (or label) is most prominent along the genome’s path in our graph model. This is done by checking for the presence of each edge from the test genome, one species at a time. For a given edge < *A, B* > in the test genome, we test the hypothesis that it belongs to species *Sp*. To do this, the vector representation of the target node *B* is bound with the vector representation of the species *Sp*. We then search for this resulting vector within node *A*’s memory by computing the cosine distance. It should be noted that a simple “close to 0” cosine distance is insufficient for confirming the presence of an edge in a node’s memory due to varying noise levels across the graph. A node’s memory is noisier if it has many bundled connections, making the reconstruction attempts more difficult. Therefore, we apply a node-dependent distance threshold that is estimated automatically for each source node and weight. This is calculated as follows: for each node *A* in our model, we query its memory for a set of *positive controls* (i.e., nodes we know it connects to) and an equal number of *negative controls* (i.e., nodes we know it does not connect to). This generates a distribution of distances for both true and false connections. We then set the specific threshold for node *A* as the 5^th^ percentile of this combined distance distribution. An edge < *A, B, Sp* > is considered reconstructed for the species *Sp* only if the query distance falls below node *A*’s unique threshold. This entire process is repeated for every possible species *Sp* ∈ {*S*_1_, *S*_2_,…, *S*_*C*_} and for every edge in the test genome *g*_*test*_. The species that accumulates the highest number of successfully reconstructed edges across the entire test genome is chosen as the final prediction.

Following the encoding strategy previously discussed, we built a graph model comprising 44,183 nodes (i.e., unique minimizers) and 612,665 edges in approximately 3 minutes using a vector dimensionality of 50,000. The entire pipeline was developed in Python 3, and the computation was performed on a server equipped with a single Intel Xeon E5-2680 v4 Central Processing Unit (CPU) (14 cores/28 threads) running @ 2.40 GHz running CentOS Linux 7. The process utilized a maximum of 24 threads and approximately 90 GB of Random Access Memory (RAM).

We performed a retraining process as previously described, converging to a final error rate of ∼5% after 6 iterations.

After training, the final graph model was evaluated on the independent test set, which contained 542 genomes (one for each species). Of these, 472 (87.08%) genomes were correctly assigned to their respective species, resulting in an overall average reconstruction rate of 44.2%.

A key feature of our method is the reconstruction rate, which serves as a powerful confidence score for each prediction. This rate measures the percentage of a test genome’s edges that were successfully found in the graph model under the predicted species label. We observed a strong correlation between this reconstruction rate and the accuracy of the prediction. Among the 472 correctly classified genomes, 162 of them showed a high degree of confidence (reconstruction rate >70%), with an average reconstruction rate of 85% (with a standard deviation at 8.4%). Conversely, the 70 misclassified genomes had a significantly average reconstruction rate of only 8.6% (with a standard deviation at 11.6%). These results are summarized in Figure 2.

**Figure 2.**
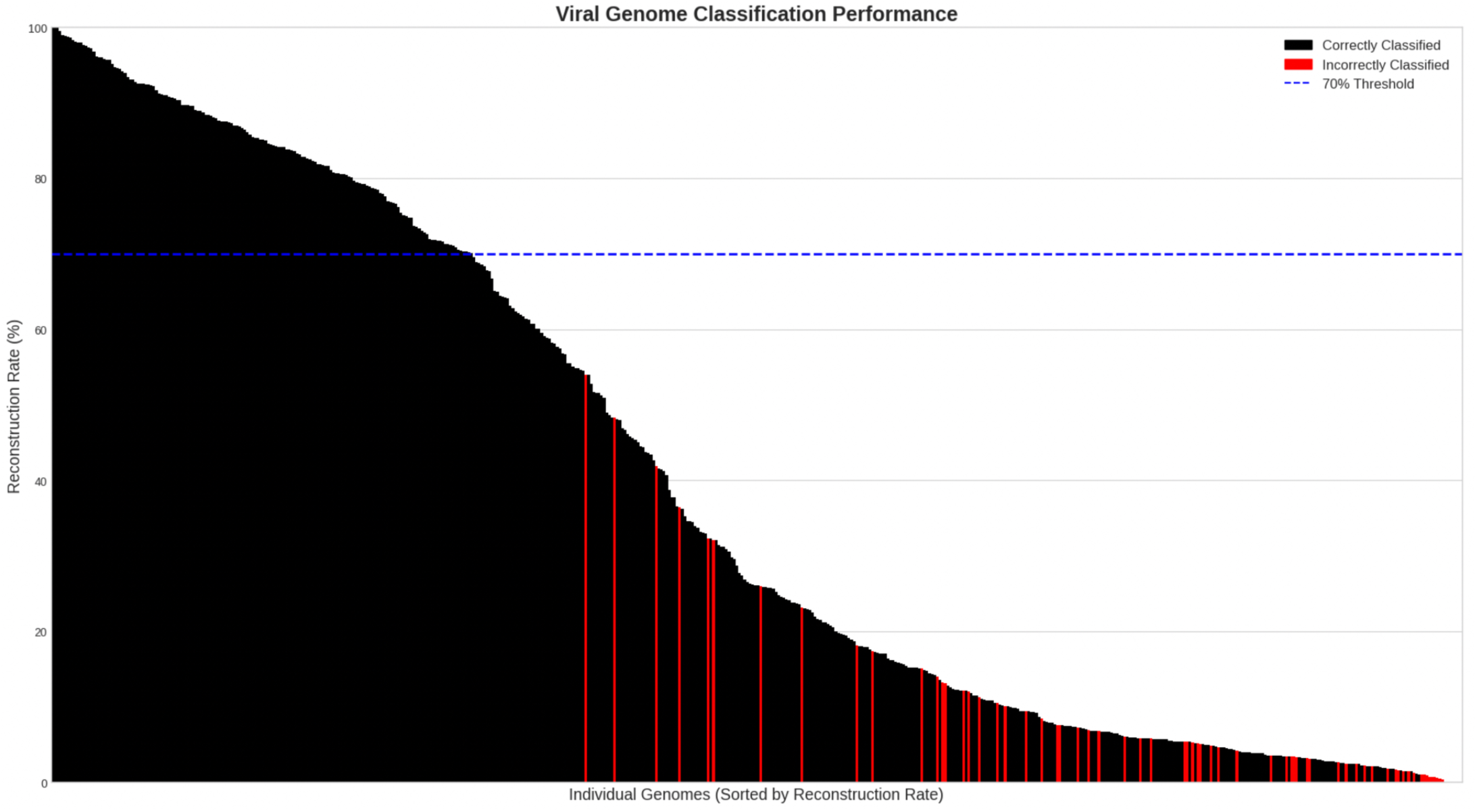
Viral genome classification performance and reconstruction rate. The figure displays the classification results of each individual test genome, sorted in descending order based on their reconstruction rate. Each vertical bar represents a single genome. Bars are colored black if the species was correctly classified, and red if it was misclassified. The dashed blue line indicates a 70% reconstruction rate threshold.

Figure 2 presents the classification performance for each test genome, showing an overview of the model’s behavior. The genomes are sorted by their reconstruction rate, which represents the percentage of the genome’s edges found within the pangenome graph. The results reveal a complex relationship between reconstruction rate, which acts as a confidence score, and classification accuracy.

The plot is dominated by a large number of correctly classified genomes (black bars). Importantly, these correct predictions span a very wide range of reconstruction rates, with many falling well below 70%. This demonstrates the model’s robustness. Even when a genome’ s edges are only partially represented in the pangenome graph (due to novel accessory genes or strain-level variation), the signal for the correct species is still significantly stronger than for any incorrect one.

This shows that the reconstruction rate functions effectively as a confidence score rather than a simple binary measure of success. A prediction with a 65% reconstruction rate can be interpreted as the model was not able to find a perfect match for the entire genome, but a significant portion of its edges point most strongly to its actual taxonomic label.

On the other hand, as expected, the frequency of misclassifications (red bars) begins to increase when getting into the low-confidence region, i.e., when the reconstruction rate drops below 50%.

Further analysis of the misclassifications revealed that they predominantly occurred between taxonomically related species despite their low reconstruction rate. It is the case of the *Gammapapillomavirus* 14 and 10 species, *Alphapapillomavirus* 9 and 11 species, and other 9 genera that were confused for one another as reported in Table 2, potentially due to their high sequence homology and shared genomic regions.

**Table 2.** Representative classification examples from the test set, illustrating the high reconstruction rates for correct predictions versus the lower rates for incorrect ones.

| Genome | True Taxonomic Unit | Predicted Taxonomic Unit | Reconstruction Rate |
| --- | --- | --- | --- |
| GCA_004288535.1 | *Gammapapillomavirus_14* (species) | *Gammapapillomavirus_10* (species) | 3.56% |
| GCA_003177875.1 | *Alphapapillomavirus_9* (species) | *Alphapapillomavirus_11* (species) | 7.57% |
| GCA_001461525.1 | *Iotapapillomavirus* (genus) | *Gammapapillomavirus* (genus) | 12.13% |
| GCA_004788075.1 | *Mischivirus* (genus) | *Megrivirus* (genus) | 4.85% |
| GCA_013087405.1 | *Potamipivirus* (genus) | *Cosavirus* (genus) | 3.31% |
| GCA_004289455.1 | *Gammapapillomavirus* (genus) | *Deltapapillomavirus* (genus) | 2.08% |
| GCA_023122835.1 | *Jeilongvirus* (genus) | *Paraavulavirus* (genus) | 8.40% |
| GCA_003178355.1 | *Betapapillomavirus* (genus) | *Alphapapillomavirus* (genus) | 4.14% |
| GCA_000892175.1 | *Betapapillomavirus* (genus) | *Alphapapillomavirus* (genus) | 3.41% |
| GCA_000973295.1 | *Genomoviridae* (genus) | *Gemycircularvirus* (genus) | 0.98% |
| GCA_002817335.1 | *Rosavirus* (genus) | *Megrivirus* (genus) | 3.38% |

A detailed breakdown of the species-level classification performance is provided in Supplementary Table S2, which lists the accession number, the ground-truth taxonomic label from NCBI GenBank, the label predicted by our model, and the corresponding reconstruction rate for each test genome.

### 3.4. The challenge of genus-level pangenome signatures

In biological taxonomy, a genus is a higher-level grouping that encompasses one or more closely related species. Moving from species-level to genus-level labels therefore reduces granularity, effectively merging related species into a broader category. Given that a majority of the species-level misclassifications occurred between closely related viruses, we hypothesized that reducing the taxonomic granularity to the genus level would resolve these ambiguities and improve overall performance. The expectation was that a genus signature would be a more robust and easier target for the model to learn.

To investigate this, we repeated the analysis by collapsing the taxonomic labels from species to genus, resulting in 218 unique genera for the classification task. Contrary to our initial hypothesis, this approach led to a significant decrease in classification performance. The model correctly classified 328 out of 542 genomes, yielding an overall accuracy of 60.51%, a notable drop from the 87.08% achieved at the species level.

The degradation in performance is visually apparent in Figure 3. Compared to the species-level results (Figure 2), the genus-level analysis shows a much larger proportion of misclassified genomes (red bars) and the rate of correct classifications falls off much more steeply. This result suggests that aggregating diverse species into a single genus label can be detrimental to the model’s performance due to signal dilution. While species within a genus are related, they can still possess significant genomic diversity, including distinct accessory genes and structural variations. By assigning the same genus label to all edges from these varied species, we effectively create a composite signal that is less coherent to find a clear, representative path for a test genome within this pangenome. This indicates that for our VSA-based approach, highly specific labels are more effective.

**Figure 3.**
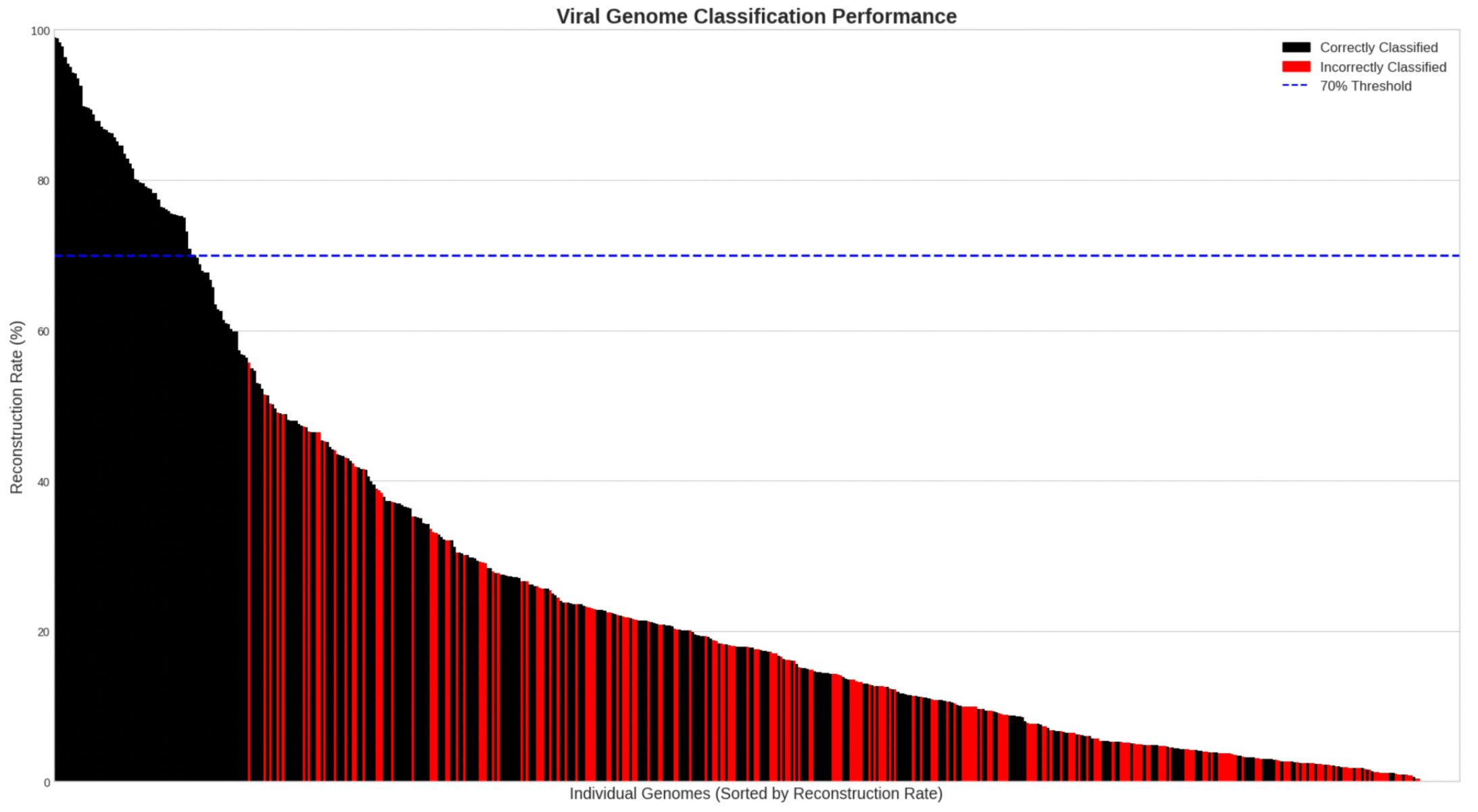
Genus-level classification performance. The figure shows the classification results aggregated at the genus level. Each bar represents a single test genome, sorted in descending order by reconstruction rate. Bars are colored black if the genome was assigned to the correct genus, and red if it was misclassified. The dashed blue line indicates the 70% reconstruction rate threshold.

Detailed results for the genus-level classification, including accession numbers, true and predicted genera, and reconstruction rates for each test genome, are provided in Supplementary Table S3.

### 3.5. Hierarchical classification with genus-level models

The previous analysis revealed a critical trade-off. The global, species-level model struggled with ambiguity between closely related species, while the genus-level model suffered from signal dilution by averaging diverse species into a single noisy label. This suggests that the problem lies not with the taxonomic rank itself, but with the “one-size-fits-all” structure of a single, monolithic pangenome graph.

We therefore hypothesized that a divide-and-conquer strategy would yield better results. By creating specialized, more focused pangenome graphs for each genus, we could resolve ambiguity without losing signal integrity. This led to a two-step hierarchical classification framework.

First, we constructed a separate pangenome graph model for each viral genus that contained multiple species. Each genus-specific graph was built exclusively from the genomes belonging to that genus but still retained the individual species labels as edge weights.

The classification of a test genome then proceeds in two stages:

1. <u>Genus prediction</u>: the test genome is queried against every genus-specific graph. The genus whose graph yields the highest reconstruction rate for the test genome’s path is selected as the predicted genus;
2. <u>Species prediction</u>: once the genus is identified, the species prediction is performed only within that genus’s specialized graph. This drastically reduces the search space and eliminates the possibility of confusion with species from other genera.

This hierarchical strategy produced a surprising and informative result. The average reconstruction rate for the initial genus prediction step increased significantly. This indicates that by querying smaller, more homologous graphs, the model was able to find much cleaner and more complete matches for the test genomes, leading to a higher degree of confidence in its assignments.

However, the higher confidence did not translate to higher accuracy. The final species-level classification accuracy decreased significantly, dropping to 33.57%. This seemingly paradoxical outcome reveals a critical bottleneck in the hierarchical approach: error propagation from the initial genus prediction step.

A second, more subtle challenge emerged even when the first step was successful. Among the genomes where the correct genus was identified with high confidence (reconstruction rate >80%), there were still eight final species-level misclassifications. As shown in Table 3, these errors exclusively occurred between extremely similar species within the correctly identified genus.

**Table 3.** Intra-genus misclassification in the high-confidence hierarchical model. The table details the eight instances where the final species prediction was incorrect, despite the model first correctly identifying the genus with a high reconstruction rate (>80%). These errors exclusively occur between similar species, highlighting the model’s final challenge in resolving ambiguity at the lowest taxonomic levels.

| Genome | True genus | True species | Predicted species | Reconstruction rate |
| --- | --- | --- | --- | --- |
| GCA_001717415.1 | *Ranavirus* | *Frog virus 3* | *European North Atlantic ranavirus* | 85.06% |
| GCA_006401535.1 | *Mastadenovirus* | *Human mastadenovirus 8* | *Human adenovirus sp* | 96.92% |
| GCA_009738835.1 | *Aviadenovirus* | *Fowl aviadenovirus D* | *Fowl adenovirus* | 84.20% |
| GCA_025023725.1 | *Aviadenovirus* | *Fowl adenovirus* | *Fowl aviadenovirus C* | 95.05% |
| GCA_004029355.1 | *Circovirus* | *Porcine circovirus 3* | *Porcine circovirus* | 96.62% |
| GCA_004086635.1 | *Circovirus* | *Porcine circovirus type 1 2a* | *European catfish circovirus* | 97.99% |
| GCA_004041475.1 | *Circovirus* | *Circovirus sp* | *European catfish circovirus* | 92.02% |
| GCA_004033655.1 | *Circovirus* | *Porcine circovirus 1* | *Duck circovirus* | 96.12% |

Additionally, the model frequently assigned a test genome to the wrong genus with high confidence (high reconstruction rate). For instance, a genome from the species *Goatpox virus* belonging to the genus *Capripoxvirus* might have a path that is ∼78% reconstructed within the *Varicellovirus* genus graph. The model would then confidently, but incorrectly, select *Varicellovirus* as the predicted genus. The subsequent species prediction, now operating within the wrong context, is doomed to fail. This demonstrates that while specialized models are powerful, their effectiveness is entirely dependent on correctly routing the query to the right model in the first place.

This experiment, despite lowering overall accuracy, provides a crucial insight: the primary challenge lies in accurately discriminating genera that share conserved genomic backbones. Results are summarized in Figure 4, while Table 4 summarizes the performance across the three different classification approaches.

**Table 4.** Comparison of classification accuracy across the three different experimental strategies with the hierarchical approach showing the average species-level accuracy and reconstruction rate.

| Classification strategy | Target level | Model structure | Overall accuracy | Average reconstruction rate |
| --- | --- | --- | --- | --- |
| Flat classification | Species | Single global graph | 87.08% | 39.60% |
| Flat classification | Genus | Single global graph | 60.51% | 25.45% |
| Hierarchical classification | Species | One graph per genus | 33.57% | 70.32% |

**Figure 4.**
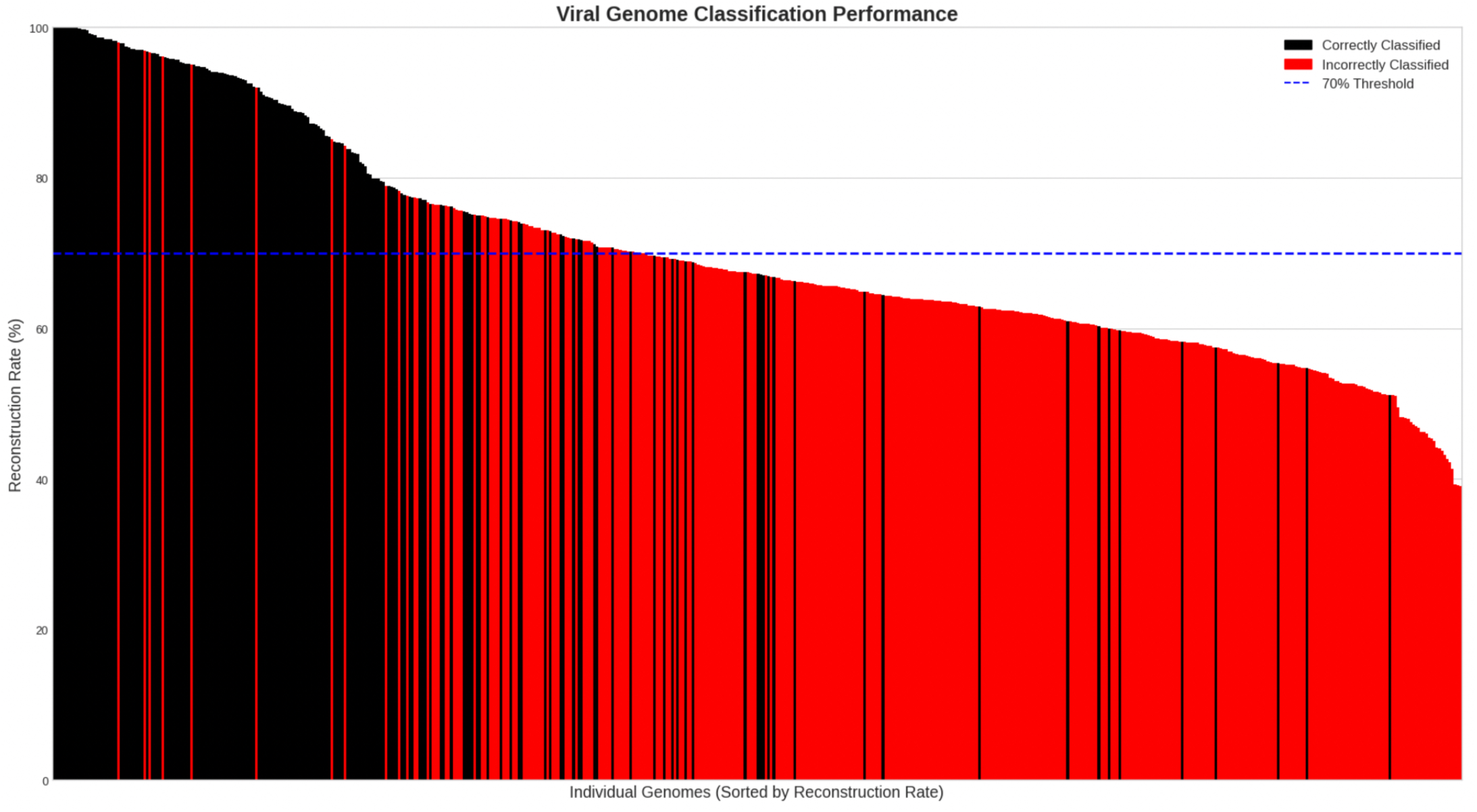
Performance of the hierarchical classification model. The figure displays the final species-level classification results from the two-step hierarchical model. Each bar represents a single test genome, sorted in descending order by the reconstruction rate achieved during the initial genus-prediction step. Bars are colored red if the final prediction was correct and black if it was misclassified. The dashed blue line indicates a 70% reconstruction rate threshold.

To provide a granular view of the hierarchical model’s performance, Supplementary Table S4 documents the outcome for each test genome. It lists the accession number, the true species label, the final predicted species, and the reconstruction rate that informed the initial genus-level assignment.

## 4. DISCUSSION AND CONCLUSIONS

This work introduces a novel approach to viral pangenome-based species classification based on a novel graph encoding technique grounded into the Hyperdimensional Computing paradigm. Pangenomes are indeed represented as weighted *de Bruijn* graphs encoded into high-dimensional vectors. We tested our method with all the viral reference genomes in NCBI GenBank, producing a vector representation of the viral pangenome graph comprising 1,084 genomes distributed over 542 species, keeping 1 genome for each species out of the training step. These last genomes have been then used to test for the accuracy of our model in retrieving genome information from the graph hypervector.

### 4.1. Model performance and the challenge of taxonomic granularity

Our results revealed a complex and counter-intuitive landscape. The most straightforward approach, a single global graph with species-level labels, yielded the highest classification accuracy at 87.08%. While this represents a strong performance for a novel method, it is crucial to place this result in the context of the current methodological landscape. A direct benchmark is challenging since our VSA-based approach represents a new paradigm for using pangenomes as a direct classification engine. The most relevant comparison is with alignment-free distance estimation tools that typically operate on the Jaccard similarity of k-mer sets [54–56]. In this context, our method explicitly encodes the graph of k-mer adjacencies, thereby retaining sequential and structural information that is discarded by set-based approaches. Therefore, our 87.08% accuracy is significant not only for its competitive performance but also for the novel, graph-centric methodology used to achieve it. It demonstrates the viability of using holographic pangenome representations as a fast and accurate alternative to both computationally intensive alignment-based methods and existing set-based sketching techniques.

This strong baseline performance makes the subsequent results even more informative. Surprisingly, attempts to simplify the problem by aggregating labels at the genus level, or to improve it with a hierarchical model, led to significantly worse performance. The flat genus-level model suffered from signal dilution, where the diversity within genera created noisy, incoherent labels, dropping the accuracy to 60.51%.

Most informatively, the hierarchical model failed due to error propagation, achieving the lowest accuracy (33.57%) despite having the highest average reconstruction rate (70.32%). This demonstrates a critical finding: the model’s reconstruction rate is a measure of its confidence, not necessarily its correctness. The hierarchical model was often “confidently wrong”, making a high-confidence but incorrect genus prediction in the first step, which doomed the subsequent species-level classification. This reveals that the primary challenge for this architecture is not resolving ambiguity between the most closely related species, but accurately discriminating between the genomic backbones of different genera.

### 4.2. Future directions

While this work presents a framework with significant potential, our results guide future directions toward solving these newly identified challenges. The immediate focus should be on improving the initial genus-prediction step of the hierarchical model, perhaps by incorporating different encoding strategies or features that are more robust to inter-genus homology. Furthermore, investigating methods to counteract signal dilution in global models could make them more effective for broader taxonomic ranks.

Additionally, expanding the pangenome to a wider range of viral species, including those present in NCBI for which no reference genomes are currently available and those yet to be discovered [57,58], is crucial for broadening the applicability of this method. In this regard, we are indeed planning to produce and maintain different models built over genomes belonging to different microbial domains. The use of minimizers to be considered over the classical definition of k-mers could also lead to more succinct vector representations of the pangenome graphs with potentially comparable classification performance. Beyond classification, we will systematically adapt and benchmark our method as a genome-assembly engine with the aim of reconstructing microbial genomes from metagenomic samples.

Ultimately, this work paves the way for targeted hardware acceleration of bioinformatics systems. Leveraging the lightweight and parallelizable arithmetic of HDC, future designs could implement system-on-chip architectures optimized for genomic graph processing. Such low-cost, energy-efficient hardware would enable scalable genomic search analyses and support the long-term vision of continuous, high-throughput genomic surveillance systems powered by dedicated HDC processors.

### 4.3. Broader impact and potential applications

The potential applications of this method extend beyond the simple classification. Integrating this approach with real-time viral surveillance systems could enable rapid identification and characterization of emerging viral threats [59]. Applying this method to outbreak analysis could provide valuable insights into the transmission dynamics and evolutionary trajectories of viruses, aiding in the development of targeted interventions [60]. Furthermore, the scalability of this approach opens up possibilities for analyzing large metagenomic datasets, potentially uncovering novel viral diversity in different environments [61]. In addition to these applications, we are releasing the full implementation of our framework as open-source software (see Availability section below). This includes both the complete pipeline for pangenome-based classification and a general-purpose HDC operation library. We aim to maximize the broader impact across multiple fields by making the source code accessible as open-source. For biomedical research, the code provides a reproducible and extendable foundation for future studies in viral genomics and metagenomics. For the hyperdimensional computing community, it offers a benchmark-ready implementation that bridges theory with practice. This joint contribution creates a “melting pot” between computational biology and emerging computing paradigms, filling an important gap between the two disciplines and opening opportunities for cross-disciplinary innovation.

### 4.4. Conclusions

Here, we introduced and evaluated a novel VSA-based framework for viral species classification. Our findings demonstrate that a global pangenome graph with species-specific labels can achieve high classification accuracy, validating the potential of this alignment-free approach. However, our most significant contribution lies in uncovering the limitations and nuances of this architecture. We show that attempts to simplify or hierarchically structure the classification problem can degrade performance due to identifiable issues or signal dilution and error propagation. This highlights a critical insight: for VSA-based pangenome analysis, the most direct and specific representation is the most effective, and the model’s reconstruction rate serves as a powerful but nuanced confidence score that must be interpreted in context. Ultimately, this study provides both a promising new tool for scalable genomics and a clear roadmap for the future research required to overcome the inherent challenges of classifying viruses.

## ADDITIONAL INFORMATION

### Availability

The whole set of genomes used for training and testing our graph models are available on NCBI GenBank. The complete list of genomes with their accession numbers and full taxonomic label is available in the Supplementary Tables S1-4.

We integrated our encoding scheme, classification algorithm, and the analysis pipeline into *hdlib* [62], an open-source Python 3 package available on the Python Package Index (PyPI – *pip install hdlib*) and Conda (*conda install -c conda-forge hdlib*). The entire pipeline is available on GitHub at https://github.com/cumbof/hdlib/tree/main/examples/pangenome.

## Abbreviations

CPU: Central Processing Unit
HDC: Hyperdimensional Computing
MAP: Multiply-Add-Permute
NCBI: National Center for Biotechnology Information
PyPI: Python Package Index
RAM: Random Access Memory
VSA: Vector-Symbolic Architecture

## Author Contribution

FC and DB conceived the research; FC and SA designed the encoding and classification model; FC implemented the software and collected the reference genomes; FC and KD built and tested the graph-based model; FC, KD, JJ, and DC performed the analysis and discussed the results; DB supervised the research; FC, KD, JJ, DC, SA, and DB wrote the manuscript and agreed with its final version.

## Acknowledgments

We would like to acknowledge the use of AI in refining the clarity and readability of this manuscript. The AI assistance was primarily used for tasks such as sentence restructuring, word choice suggestions, and identifying potentially unclear phrasing. We emphasize that the AI was used solely for language enhancement and did not contribute to the generation of research ideas, data analysis, code implementation, or the interpretation of results. All conclusions drawn and insights presented in this manuscript are solely the product of the authors’ own analysis and expertise.

